# Bees can be trained to identify SARS-CoV-2 infected samples

**DOI:** 10.1101/2021.10.18.464814

**Authors:** Evangelos Kontos, Aria Samimi, Renate W. Hakze-van der Honing, Jan Priem, Aurore Avarguès-Weber, Alexander Haverkamp, Marcel Dicke, Jose L Gonzales, Wim H.M. van der Poel

## Abstract

The COVID-19 pandemic has illustrated the need for the development of fast and reliable testing methods for novel, zoonotic, viral diseases in both humans and animals. Pathologies lead to detectable changes in the Volatile Organic Compound (VOC) profile of animals, which can be monitored, thus allowing the development of a rapid VOC-based test. In the current study, we successfully trained honeybees (*Apis mellifera*) to identify SARS-CoV-2 infected minks (*Neovison vison*) thanks to Pavlovian conditioning protocols. The bees can be quickly conditioned to respond specifically to infected mink’s odours and could therefore be part of a wider SARS-CoV-2 diagnostic system. We tested two different training protocols to evaluate their performance in terms of learning rate, accuracy and memory retention. We designed a non-invasive rapid test in which multiple bees are tested in parallel on the same samples. This provided reliable results regarding a subject’s health status. Using the data from the training experiments, we simulated a diagnostic evaluation trial to predict the potential efficacy of our diagnostic test, which yielded a diagnostic sensitivity of 92% and specificity of 86%. We suggest that a honeybee-based diagnostics can offer a reliable and rapid test that provides a readily available, low-input addition to the currently available testing methods. A honeybee-based diagnostic test might be particularly relevant for remote and developing communities that lack the resources and infrastructure required for mainstream testing methods.

## Introduction

Infections and other pathologies lead to physiological changes in the bodies of animals (Trabue *et al*., 2010) and humans (Buljubasic & Buchbauer, 2015; Sethi *et al*., 2013; Shirasu & Touhara, 2011, Probert *et al*., 2009). Consequently, the emitted volatile organic compounds (VOCs) differ between healthy and infected individuals (Fitzgerald *et al*., 2017; Wilson *et al*., 2018; Olsson *et al*., 2014; Trabue *et al*., 2010; Probert *et al*., 2009). VOCs constitute an odour fingerprint depending on age, sex, diet, genetic background, and metabolic conditions, thus making this odour fingerprint unique for every individual (Buljubasic & Buchbauer, 2015; Shirasu & Touhara, 2011). Analysing that fingerprint can provide relevant information about the state of the individual’s health. VOC analysis has been consequently used for disease diagnostics, mostly in the form of breath and faeces analysis in both humans and animals (Fitzgerald *et al*., 2017; Wilson *et al*., 2018; Olsson *et al*., 2014; Trabue *et al*., 2010; Probert *et al*., 2009).

The current COVID-19 pandemic has clearly shown the need for both the rapid development of diagnostic tests and the rapid delivery of reliable results (European Centre for Disease Prevention and Control, 09/2020). Fast and reliable diagnostic tests are required to effectively implement control measures such as quarantine of infected people or animals (Wells *et al*., 2021). There is a global need for reliable and rapid testing, which has led to the development of very reliable PCR tests and rapid SARS-CoV-2 tests such as the RNA RT-LAMP (Fowler *et al*., 2021) and antigen tests (Krüttgen *et al*., 2021). However, in developing countries and remote areas such methods may not be easily available. Dogs have been successfully trained to discriminate between SARS-CoV-2-infected and non-infected individuals with a diagnostic sensitivity ranging from 65% to 82.6% and specificity of 89% and 96.4% respectively (Eskandari *et al*., 2021; Jendrny *et al*., 2020). Similar to dogs, some insects have keen olfactory capabilities. For example, fruit flies (*Drosophila melanogaster*) can detect cancer in humans (Strauch *et al*., 2014), while honeybees (*Apis mellifera*) have exhibited the ability to detect some human diseases, such as tuberculosis (Suckling & Sagar, 2011). Honeybees can, therefore, be a potential alternative to dogs for the detection of COVID-19 with the benefit of being readily available and having low costs of maintenance.

Pavlovian conditioning was first applied to dogs (Pavlov, 1927) and later to honeybees (Takeda, 1961). Bees possess the reflex to extend their proboscis when detecting a sugar solution (PER; proboscis extension reflex) and they can be conditioned to exhibit a PER when exposed to specific odours. Takeda’s (1961) classical conditioning pairs a conditioning stimulus (CS), such as an odour, with an unconditioned stimulus (US), the food reward, which in most cases is a sugar water solution (Matsumoto *et al*., 2012, Sutherland *et al*., 2010, Wright *et al*., 2010). After such training the bees exhibit PER when exposed to the CS, without the presence of sugar water.

Previous studies have shown that animals can detect differences between VOCs emitted by healthy or SARS-CoV-2 infected individual animals or humans (Eskandari *et al*., 2021; Jendrny *et al*., 2020; Suckling & Sagar, 2011). The objective of this study was to assess the potential of training bees for the detection of SARS-CoV2-infected animal samples. We assessed two different training methods and show that bees can be effectively trained to detect differences in odours between samples collected from SARS-CoV2 infected and uninfected minks (*Neovison vison*), highlighting the potential of a honeybee-based diagnostic test for the detection of diseases.

## Materials and Methods

### Honeybees’ preparation

At the start of each experimental day during April and May 2021, we collected a new batch of honeybees (*Apis mellifera*) from the same beehive, located 2 km away from the Wageningen Bioveterinary Research (WBVR) laboratory in Lelystad, the Netherlands. We assumed that the bees were a mixture of different working classes. Foragers were preferred but the weather conditions did not allow for flights every day so discrimination between worker classes was not always possible. The bees were collected with a brush from inside the hive or by collecting departing bees at the hive entrance, using the same brush. For transport to the laboratory, bees were placed in transparent cylindrical plastic containers (100 ml), that carried 5-15 bees each. A total of 149 bees were used during the experiments.

The containers with honeybee workers were placed in a freezer (−20°C) for 3-5 min until the bees become inactive, which makes harnessing safer. Once out of the freezer, the bees were placed on a paper towel and they were inserted inside our custom-made “bee-holders” with the help of tweezers (Fig. 1). The bee-holders are made of plastic and have the following dimensions: 20 × 10 × 10 mm. They consist of two parts, the back and base which allows the experimenter to hold it easily and the front part, the chamber, where the bee is kept. The chamber has two openings, one in the bottom to allow for the bee to be inserted easily and a door-like structure above. The door closes once the bee is inside the chamber locking its head into position, while allowing the rest of the body to move freely. The chamber also has two openings for the bee’s wings, avoiding unnecessary injuries. We harnessed the bees 30 min after collection and the experiments started 3 h after harnessing. We collected and harnessed multiple bees in parallel. Those that exhibited a Proboscis Extension Reaction (PER) after a brief touch of the antenna with the sugar-water solution (Fig. 1), were used for conditioning.

**Figure 1.**
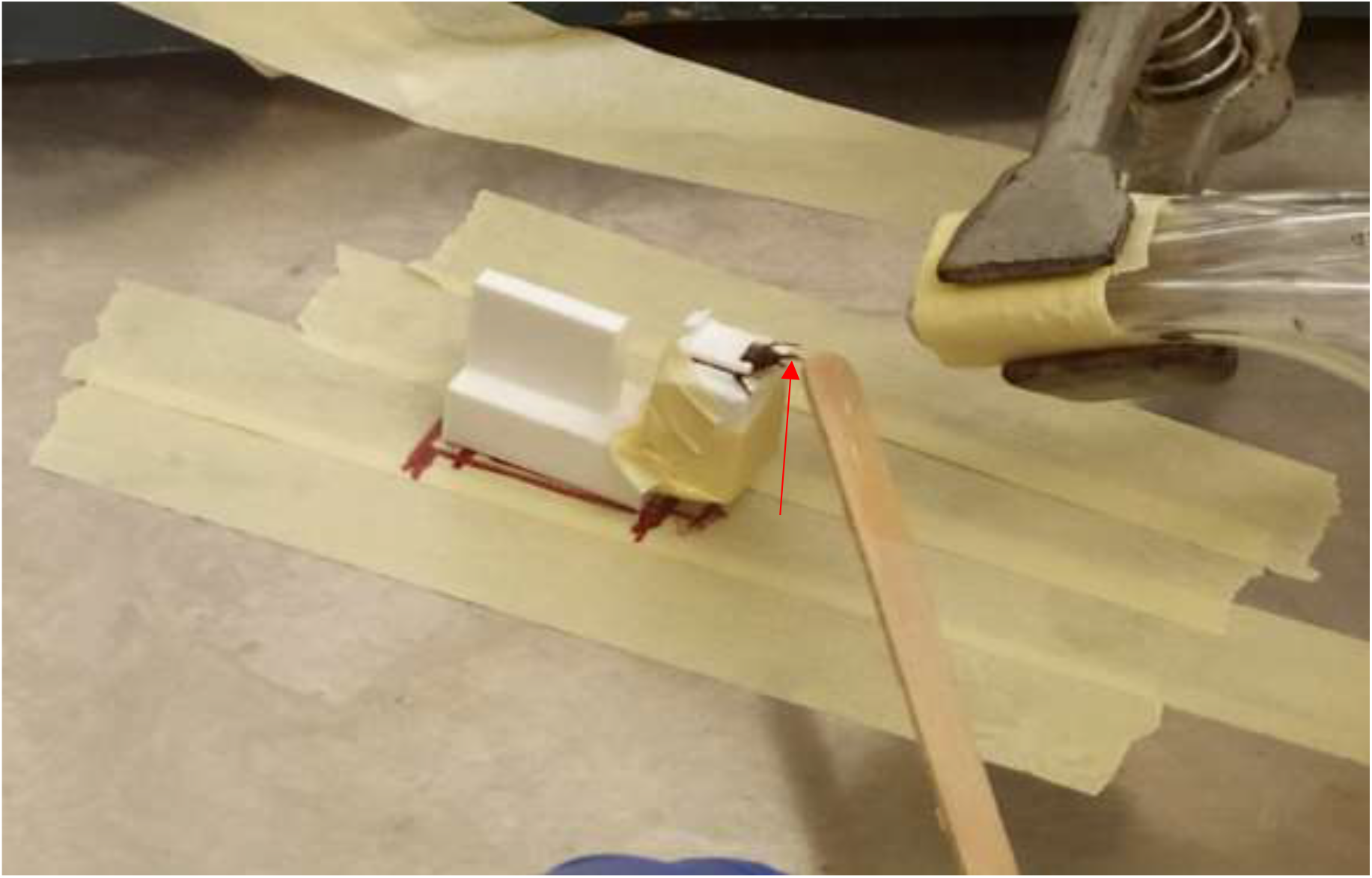
Picture of the conditioning procedure during protocol 1. A single honeybee harnessed inside our custom-made bee-holder. The bee has just been exposed to a positive sample and been provided with a wooden stick soaked in sugar water, which has led her to express the Proboscis Extension Reflex (PER). The red arrow indicates the bee’s proboscis.

### Sample selection

Throat swabs were taken of necropsied animals from a mink farm during the SARS-CoV2 epidemic in the period of April-November 2020, in the Netherlands. 2 ml of Dulbecco’s Modified Eagle Medium (DMEM), supplemented with 10% Fetal Calf Serum (FCS) and 1% Antibiotic Antimycotic (Gibco, Thermofisher, Netherlands) was added to each swab sample. The presence of viral SARS-CoV-2 RNA as well as the Cycle Threshold (Ct) value of the samples were determined by real time RT-PCR on the SARS-CoV-2 E gene (Corman *et al*., 2020). All minks were fed the same feed ration and were raised under the same conditions in the same location in a production farm in the South of the Netherlands (Oreshkova *et al*., 2020). The swab solutions (60 μl of liquid) from SARS-CoV-2 infected and healthy minks were absorbed by filter papers (Whatman, Cat No 1001090) (1 × 3 cm size), which were placed inside identical syringes (20 ml) and the plastic containers.

### Olfactory conditioning procedures

We tested two different bee training protocols inspired by earlier research reported by Sutherland *et al*. (2010) (Protocol 1) and Wright *et al*. (2010) (Protocol 2).

All experiments were executed in a biosafety level 2+ laboratory at WBVR, in Lelystad, the Netherlands. The bee conditioning and retention test took place inside a biosafety cabinet. The bees were introduced inside the biosafety cabinet after being harnesses and remained inside until the end of the experiment. The airflow inside the hood was 0.36 m/s, the temperature 21 °C and the humidity 56%; these conditions were regulated throughout the experiment. A trial lasted 40 seconds during which the bee was placed in front of the odour delivery apparatus. The syringes released an air puff after the first 20 s, that lasted for 5 s, during which we recorded the bee’s reaction. The bee would stay there for 15 s and would then be replaced by the next bee in line. The ITI (Intertrial Interval) was 10 min in both protocols.

### Unconditioned stimuli

We used two different protocols, in both of which, a wooden stick was soaked with sugar water (US) (Fig. 1), first touching the bee’s antennae, to induce PER, and later the proboscis. If the proboscis was already extended, the antennae were not touched. The sugar water reward occurred for 5 s with a 2 s overlap with the air puff from the syringe, which preceded it. The US used during protocol 1 was 1.5 M sugar-water solution (Sutherland *et al*., 2010). In protocol 2 we used two US types. The positive unconditioned stimulus (US+) was a 1 M sugar-water solution. During protocol 2, we also exposed the bees to a quinine-sugar-water solution, an aversive stimulus, the negative US (US-; 300 mM sugar, 10 mM quinine; Wright *et al*., 2010). When the bees were exposed to samples from healthy mink individuals, the sugar-soaked stick first touched the antennae to induce PER, and then the quinine-sugar-soaked stick would touch the proboscis. If the proboscis was already extended the antennae were not touched.

#### Protocol 1

In this training procedure we used one sample from an infected (positive), and one from a healthy (negative) animal to condition 56 honeybees. The bees were trained with the same positive sample, each experimental day, for which a standardized cycle threshold (Ct) value of 21 was acquired from a PCR test. The filter paper soaked with the negative sample was placed inside a small plastic container connected to two tubes. One tube was connected to a pump, providing a constant air flow (40 ml/min) while the other tube was placed in front of the bees, thus delivering the healthy sample odour constantly during the training trials. The syringe containing the infected sample was connected with a similar tube. The air flow necessary to deliver the infected sample odour to the bees was provided by manually operating the syringe. The tubes from the plastic container (healthy sample) and the syringe (infected sample) were taped together, so that the bee could be exposed to both simultaneously during the CS delivery time. There was a distance of 2 cm between the bees and the tube outlets and the syringe released an air puff of 15 ml in 5 s.

We performed: i) seven conditioning trials in which the bees were exposed to the positive infected sample against the background of the healthy negative sample and were provided with a sugar-water solution as unconditioned stimulus (US); and ii) 7 trials in which the bees were only exposed to the healthy sample and no US. The trials followed a pseudorandomized order (H-I-H-I-I-H-I-H-H-I-I-H-H-I) (H: Healthy, I: Infected) (Matsumoto *et al*., 2012). During conditioning we recorded the number of bees that expressed PER during each of the seven training rounds, before exposure to US, to assess the rate with which they learned (learning curve).

#### Protocol 2

In this training procedure we used three samples from infected animals (positive) and three samples from healthy ones (negative) to condition 92 honeybees. The bees were trained with positive samples for which a standardized Ct value (21) had been recorded in the PCR and tested with negative samples with three different Ct values (21, 27, 30). The filter papers containing the samples were placed inside identical syringes and were placed in front of the bees. There was a distance of 2 cm between the bees and the syringe outlets, which released an air puff of 20 ml in 5s.

We performed nine conditioning trials in which the bees were exposed to the positive samples and nine trials in which the bees were exposed to the negative samples (three trials for each sample). The bees were given the US+ when exposed to positive (infected) samples and the US-when exposed to negative (healthy) ones. The trials followed a pseudorandomized order (H-I-H-I-I-H-I-H-H-I-I-H-H-I-H-I-I-H) (H: Healthy, I: Infected) (Matsumoto *et al*., 2012). The different samples were also randomised as follows (A-B-C-C-B-A-B-A-C). In addition, the experiments were mirrored, so that half of the bees would be exposed to exactly the inverse of the (H-I-H-I-I-H-I-H-H-I-I-H-H-I-H-I-I-H) and (A-B-C-C-B-A-B-A-C) order. As a result, half of the bees started with a sugar reward (infected mink samples) and finished with healthy mink samples (quinine punishment) and the other half followed the reverse order. By comparing between these sequences (punishment first, reward first), we analysed which one yields the best results. Neither sequence yielded significantly better results during 1 h, while during the 24 h retention only one comparison was significantly different (Supplementary Fig. 1). This indicates that the sequence with which the samples are provided to the bees does not significantly affect their training outcome. During conditioning we recorded their learning curve and later analysed their memory retention.

### Testing: Memory retention

In both protocols we performed memory retention tests after 1 and 24h. The number of bees differed between the training phase, the 1 h retention test and the 24 h retention test, as a result of bee mortality.

#### Protocol 1

One hour after the end of the training, we performed a retention test to check the bees’ memory by exposing them to positive and negative samples without any US and recorded whether they extended their proboscis. For the retention test we neither changed the layout used during training, nor did we remove the background negative sample odour (Old healthy sample: Old-healthy). However, we introduced novel odours of a different infected mink’s swab (New infected sample: New-infected) and a different healthy mink’s swab (New healthy sample: New-healthy) and an empty syringe to test the effect of the additional air pressure. The empty syringe test was also testing the bees’ reaction to Old-healthy (which was present on the background). During the retention test at 24 h after the end of the training, no bees reacted to the empty syringe indicating that air pressure does not influence their reaction. At the same time, it confirmed that the bees had successfully been trained to ignore Old-healthy. As a result, we did not use the empty syringe during the following days in order to avoid over-testing the bees risking a dissociation between CS and US.

#### Protocol 2

One hour after the end of the training, we tested the bees’ retention capabilities. Every bee was tested multiple times with six odours in total. Two new negative and two new positive samples were used and were grouped together, during data analysis, as New-healthy and New-infected, for a more comprehensive presentation of the results. We also used the positive sample that the bees reacted to the most during conditioning (Old-infected) and the negative that they reacted to the least (Old-healthy). For the retention test we did not change the training layout and we presented the samples in a random order.

### Data analysis

We analysed the learning rate of the bees for each protocol independently by performing generalized linear mixed models (GLMM) with a binomial distribution. In these models the bees’ response (PER: 0 or 1) was set as the dependent variable, while the sample (positive or negative), the conditioning round and the interaction between samples and conditioning round were fitted as fixed explanatory variables. The bees’ individual identification was introduced as random intercept to account for multiple measures being made with each bee. Significance of the explanatory variables was assessed using the Wald test, with threshold for significance set to p < 0.05. Using these models, we were interested in assessing the improvement in the discrimination ability of bees between infected and healthy samples as a function of the number of conditioning rounds.

To assess the bee’s discrimination accuracy between healthy and infected samples at 1 h and 24 h after conditioning, we fitted again a GLMM with a binomial distribution. Models were fitted for each training protocol and for each retention time (1h or 24h) independently. In these models the bees’ response (PER: 0 or 1) was the dependent variable, while the type of sample (New-healthy, Old-healthy, New-infected, Old-infected) was the fixed explanatory variable and the bees’ identification was introduced as random intercept. For statistical comparison between sample types, we used the New-healthy sample as reference. This sample was used as reference because we considered that if the bees were to be used for diagnostic purpose they will be exposed to unknown (new) samples which they need to classify (discriminate between) as healthy or infected. Significance of the explanatory variables was assessed using the Wald test.

To explore the diagnostic potential of the bees and predict the diagnostic performance of the practical application of using bees for diagnosis of SARS-COV-2 we:

1st) Tested the association between the sample’s Ct value (indicator of virus concentration in the sample) and detection rates after 1 hr of retention. Infected samples used had Cts of 21, 27 and 30. The proportions of bees reacting to each of these samples were compared using a Chi square test. For this analysis, independence was assumed and a Bonferroni correction for multiple comparisons was applied for the interpretation of significance.

2nd) Simulated a population of infected and non-infected samples which were individually tested by a group of bees. This simulation was done by random sampling with replacement groups of 10 bees, which would be part of a diagnostic group, from the retention tests done at 1 and 24 h. Sampling was done for positive samples or negative samples independently. A total of 300 groups of 10 bees exposed to positive samples and 300 groups exposed to negative samples were simulated. Sampling was done from the dataset with the retention results, which contained diagnostic results at individual bee level. From each sampled group the number of bees preforming a correct discrimination of the sample (either positive or negative) was recorded. This number was then used to perform a Receiver Operating characteristics (ROC) analysis to identify a potential diagnostic threshold and assess the diagnostic efficacy (Sensitivity and Specificity) of the system (groups of bees).

All data analyses were performed using the statistical software R version 4.0.2 (R, 2013). The GLMM were fit using the library lme4 (Bates *et al*., 2015). ROC analyses were done using the libraries pROC and ROCR (Robin *et al*., 2011, Sing *et al*., 2005).

## Results

### Protocol 1

We analysed the bees’ learning curve during conditioning by fitting a GLMM. A significant interaction (log-odds = −0.38, standard error (SE) = 0.11, Z = −3.37, *P* < 0.001) between treatment and conditioning round was observed, which suggests a significant increase in the bee’s ability to discriminate between infected and negative samples with increasing conditioning rounds (Table 1). By the end of the conditioning phase (round 7), 37 bees out of the total 56 bees (66.1%) expressed PER towards the infected sample (Old-infected) and 4 out of 56 (7.1%) towards the healthy sample (Old-Healthy) (Fig. 2A).

**Figure 2.**
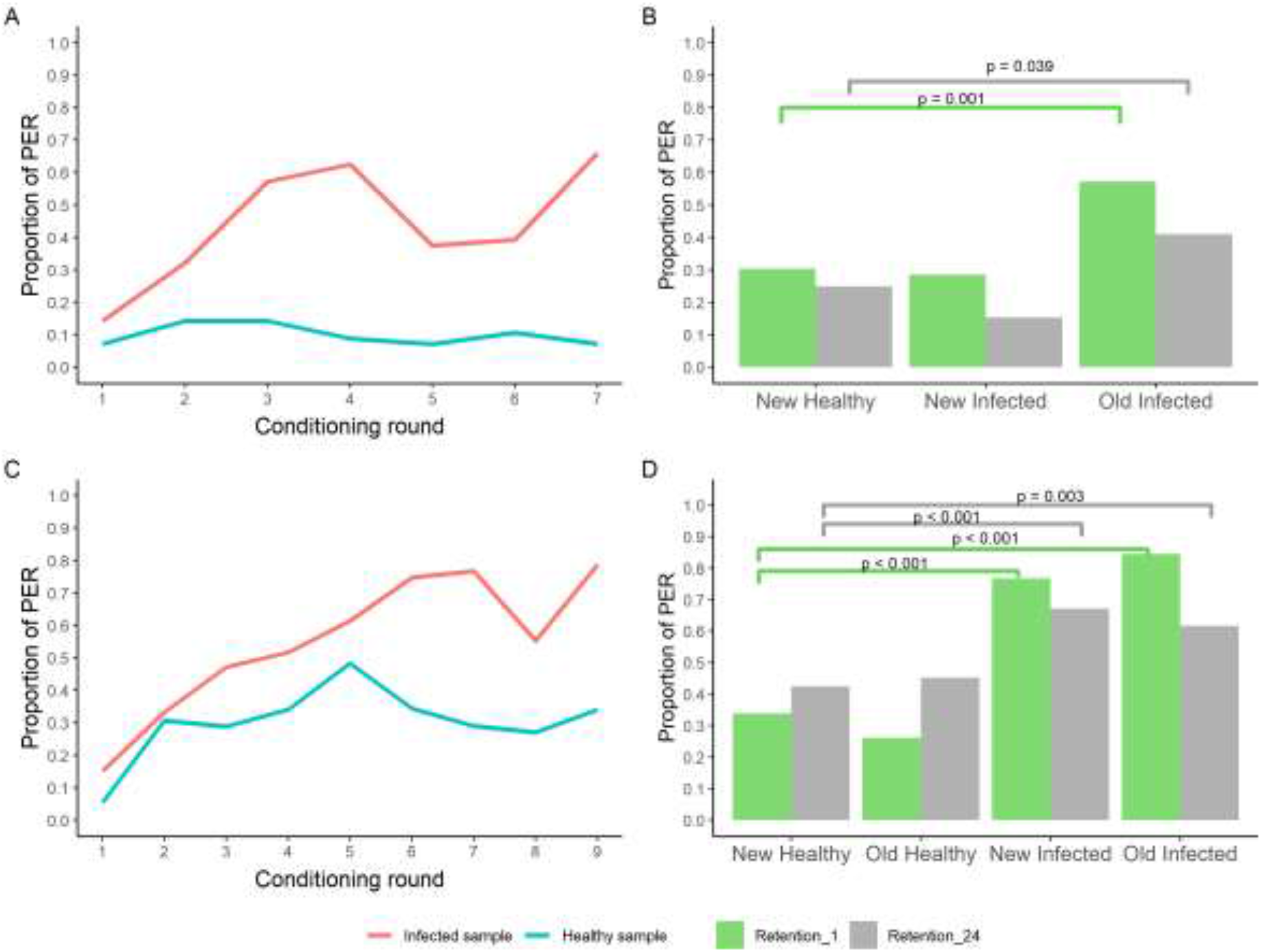
Learning and memory retention of the bees. *P*anels A and B show the learning curve (n = 56 bees) and memory retention (n = 56 bees) of bees subjected to protocol 1. Panel C and D show the learning curve (n = 92 bees) and memory retention (n = 56 bees) of bees subjected to protocol 2. For Panel A and C (learning curves), the Y-axis shows the proportion of bees expressing PER towards infected (red) and healthy (blue) samples in each conditioning round while the X-axis indicates the conditioning round. For panels C and D (memory retention, the Y-axes show the proportion of bees expressing PER and the X-axes show the different types of samples that the bees were exposed to 1 h (green columns) and 24 h (grey columns) after the conditioning training ended. Segments and corresponding *P* values indicate comparisons where significant. The sample type New Healthy was used as reference for statistical comparison.

**Table 1.**
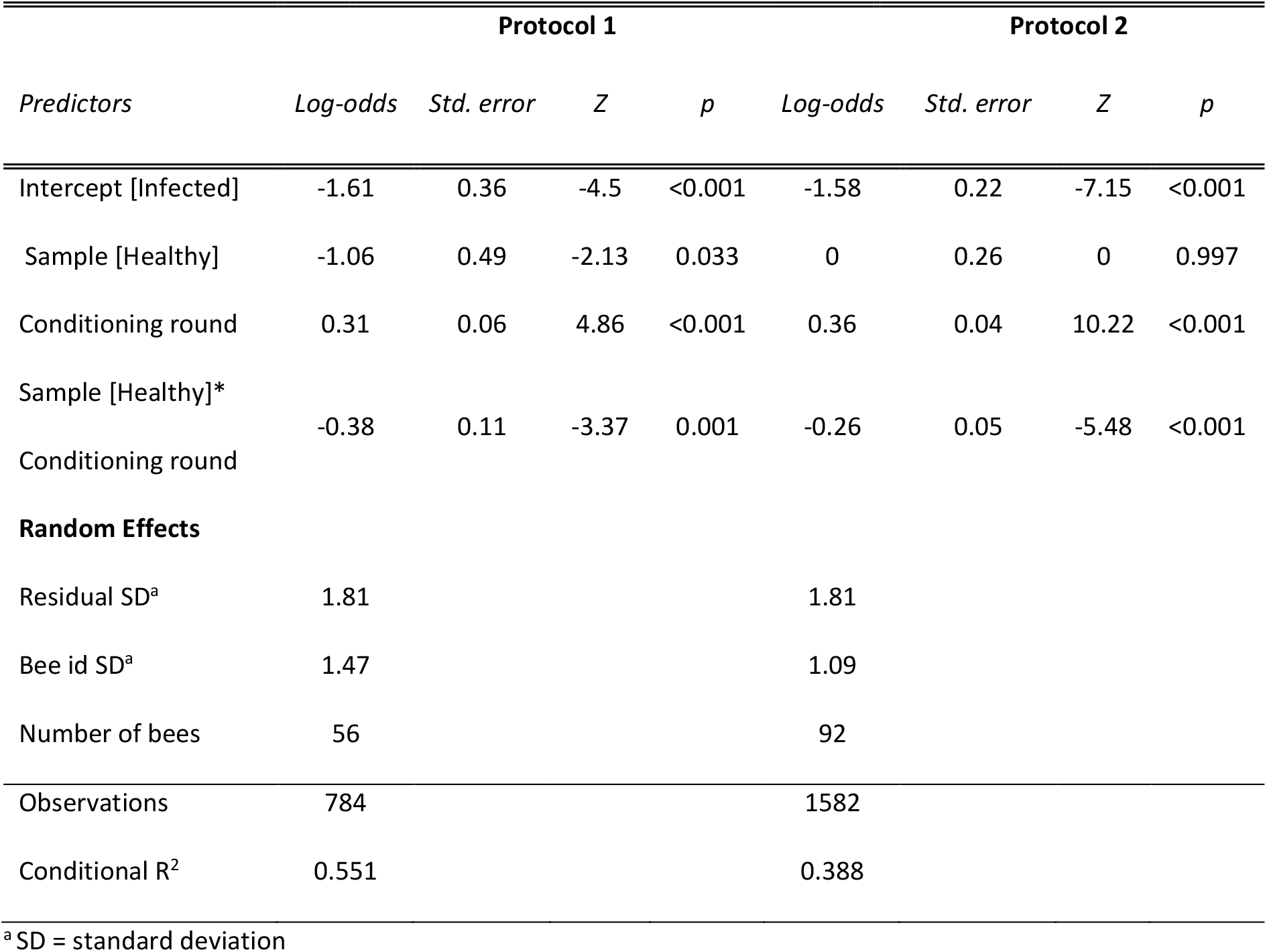
Logistic regression mixed model results analysing the bees’ learning curves

| *Predictors* | **Protocol 1** |  |  |  | **Protocol 2** |  |  |  |
| --- | --- | --- | --- | --- | --- | --- | --- | --- |
|  | *Log-odds* | *Std. error* | *Z* | *p* | *Log-odds* | *Std. error* | *Z* | *p* |
| Intercept [Infected] | -1.61 | 0.36 | -4.5 | <0.001 | -1.58 | 0.22 | -7.15 | <0.001 |
| Sample [Healthy] | -1.06 | 0.49 | -2.13 | 0.033 | 0 | 0.26 | 0 | 0.997 |
| Conditioning round | 0.31 | 0.06 | 4.86 | <0.001 | 0.36 | 0.04 | 10.22 | <0.001 |
| Sample [Healthy]*<br>Conditioning round | -0.38 | 0.11 | -3.37 | 0.001 | -0.26 | 0.05 | -5.48 | <0.001 |
| **Random Effects** |  |  |  |  |  |  |  |  |
| Residual SD<sup>a</sup> | 1.81 |  |  |  | 1.81 |  |  |  |
| Bee id SD<sup>a</sup> | 1.47 |  |  |  | 1.09 |  |  |  |
| Number of bees | 56 |  |  |  | 92 |  |  |  |
| Observations | 784 |  |  |  | 1582 |  |  |  |
| Conditional R<sup>2</sup> | 0.551 |  |  |  | 0.388 |  |  |  |

<sup>a</sup>SD = standard deviation

We tested the bees’ memory retention 1 h after the conditioning phase. Every bee was exposed to three odours (samples): Old-infected (sample used for conditioning), New-infected, New-healthy. During exposure, the Old-healthy sample was always present as a background odour. Using the New-healthy sample as reference for comparisons, the GLMM analysis confirmed that most of the bees were able to discriminate (log-odds = 1.8, SE = 0.56, Z =3.24, *P* = 0.001) the Old-infected sample (32 out of 56 (57.1% reacted to Old-infected)) from the New-healthy sample (17 out of 56 (30.4%) reacted to New-healthy). However, bees were not able to discriminate (log-odds = −0.13, SE = 0.52, Z = −0.26, *P* = 0.796) the New-infected sample (16 out 56 (28.6%)) from the New-Healthy sample (Fig. 2B).

Another retention test took place 24 h after the end of training, using the same odours we had used for the 1 h retention test. When analysing the reaction of the bees 24 h after conditioning, out of 56 bees, 14 (25%) reacted to New-healthy sample, 23 reacted to Old-infected (41.1%) sample and 8 out of 52 (15.4%%) reacted to the New-infected sample. Bees were only able to significantly discriminate between the Old-infected (Log-odds = 1.06, SE = 0.51, Z = 2.07, *P* = 0.038) and the New-healthy samples (Fig. 2B).

### Protocol 2

Similar to protocol 1, we analysed the bees’ learning curve during conditioning using a GLMM. A significant interaction (log-odds = −0.38, SE = 0.11, Z = −3.37, *P* < 0.001) between treatment and conditioning round was observed, which suggests a significant increase in the bee’s discrimination ability between the positive and negative samples with the conditioning rounds (Table 1). By the end of conditioning (round 9) 67 out of 85 (78.8 %) bees expressed PER towards infected samples and 23 out of 85 (27.1 %) towards healthy samples (Fig. 2C).

We tested the bees’ memory retention 1 h after the end of the conditioning. Overall, 71 bees out of 84 (84.5 %) reacted to the Old-infected sample. Bees reacted 129 times out of 168 trials (77.8%) to the New-infected samples while 22 bees out of 84 (26.2%) reacted to the Old-healthy samples. Bees reacted 57 times out of 168 trials (33.9%) to New-healthy samples. The GLMM analysis (using New-healthy as reference for comparison) confirmed that bees were able to significantly discriminate (express PER) between the New-infected sample and both the Old- (log-odds = 3.02, SE = 0.42, Z = 7.27, P < 0.001) and New-infected (log-odds = 2.41, SE = 0.31, Z = 7.79, P < 0.001) samples. No differences were observed between the bees’ reaction towards the Old- (log-odds = −0.47, SE = 0.33, Z = −1.40, *P* = 0.161) and New-healthy samples (Fig. 2D).

Another retention test was executed 24 h after the end of training, using the same odours we had used for the 1 h retention test. Overall, 45 bees out of 73 (61.6%) reacted to the Old-infected sample and 98 times out of 146 trials (67.1%) to the New-infected samples, while 33 bees out of 73 (45.2%) reacted to the Old-healthy samples and 62 times out of 146 trials (42.5%) to New-healthy samples. The GLMM analysis showed that bees were able to significantly discriminate between the New-infected and both the Old-infected (log-odds = 1.01, SE = 0.34, Z = 3.02, *P* = 0.003) and New-infected (log-odds = 1.31, SE = 0.28, Z = 4.59, *P* < 0.001) samples. The bees’ reaction to the Old- and New-healthy samples did not differ significantly (log-odds = 0.14, SE = 0.33, Z = 0.44, *P* = 0.658) (Fig. 2D).

### Bees as a diagnostic tool

#### Association between Ct values and the bees’ retention ability

We compared the proportion of bees showing PER depending on the Ct values of the infected samples. After being trained on a sample with a Ct =21, a total of 42 bees were exposed to samples with three different Ct values (Ct: 21, 27 or 30): 35 (83.3%) bees showed PER for the sample with a Ct = 21, 40 (95.2%) showed the samples with Ct = 27 and 31 (73.8%). The samples used for training were those with a Ct =21, hence we used this group as reference for comparison. No significant differences were observed between either the Ct = 21 group and the Ct = 27 group (*X^2^* = 0.63, df = 1, P = 0.16, Padjusted = 0.475) or Ct =21 and Ct = 30 (*X^2^* = 0.63, df = 1, P = 0.42, Padjusted = 1) groups, indicating that bees trained with samples having a high virus concentration (a low Ct value) are still able to recognize samples with lower virus concentrations (high Ct value).

#### Predicted performance when using bees as a diagnostic tool

The distribution of diagnostic results when testing healthy and infected samples in a simulated scenario in which a group of 10 trained bees would be used to test a sample, is shown in Fig. 5a. The ROC analysis on the simulated data resulted in an estimated area under the curve (AUC) (Fig. 5b) equal to 0.96 (95% confidence interval (CI): 0.95 – 0.98), which indicates that using groups of trained bees could be a diagnostic tool with significant discrimination accuracy (AUC > 0.5, p < 0.001). Using a response of 6 or more (out of 10) bees showing PER per test to classify a sample as positive would maximize the diagnostic performance of this tool. The resulting potential sensitivity (true positive rate), which is the probability that the test will correctly classify a truly infected sample as positive, would be 0.92 (95% CI: 0.89 – 0.95) and the potential specificity (true negative rate), which is the probability that the test will correctly classify a healthy sample as negative, 0.86 (95% CI: 0.82 – 0.90) (Fig. 3).

**Figure 3.**
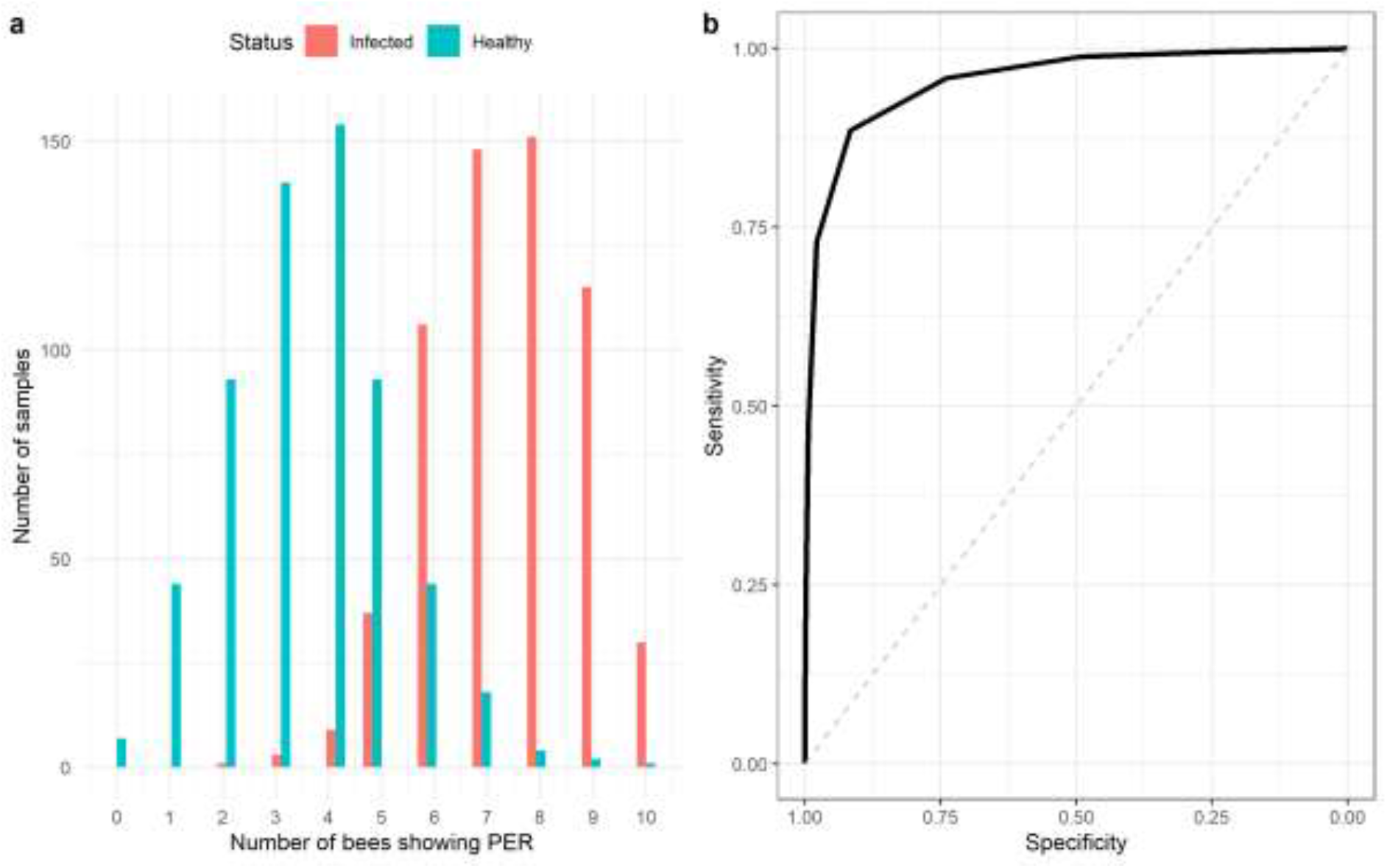
Simulated diagnostic potential of trained bees. a) Distribution of simulated diagnostic results where a group of 10 bees is used as diagnostic tool per sample. The X axis indicates the number of bees (out of 10) per test showing PER. b) Receiver operating characteristic curve (ROC) of the predicted diagnostic performance.

## Discussion

The objective of this study was to investigate whether bees can be trained to detect SARS-CoV-2 infected samples. Our data show that the differences in odour between SARS-CoV-2 infected samples and uninfected samples can be recognised by honeybees. The bees discriminated between samples taken from healthy and SARS-CoV2 infected individuals. Although the bees’ discrimination ability decreases between 1 h and 24 h post conditioning, we observed that they were nevertheless still able to significantly discriminate between new infected and healthy samples after one day post-conditioning. Moreover, their ability to recognise a positive sample was not compromised by the samples’ viral load (expressed in Ct values), since bees recognised samples with higher Ct values (lower viral load) equally well as they did with samples with low Ct values used for conditioning. By performing simulations of the potential clinical application of the bees as a diagnostic tool, we predict that bees could be effective for diagnostics with a predicted sensitivity around 92% and specificity around 86%.

With our protocol 1, we followed a similar procedure as Sutherland *et al*. (2010). In our study, more bees learned to recognize the rewarded odour (current study: 66.1%, Sutherland *et al.:* 30-40%). Sutherland *et al*. (2010) reported that one hour after training, 20% of the trained bees were no longer able to discriminate, which is similar to the reduction in percentage we observed. Sutherland *et al*. (2010) did not perform any test with novel infected and healthy samples, and we did not find literature where a similar approach to ours was taken. In protocol 2, we followed a similar procedure as Wright *et al*. (2010). The bees in the present experiment learned slightly less well than in their experiment (current study: 78.8%, Wright *et al*.:>85%). Assuming that this difference is significant, we could speculate that this is a result of samples being more difficult to discriminate either by being more perceptually similar or less concentrated. Wright *et al*. (2010) did not test the 1 h memory of the bees but only their memory after 10 min. Their results were similar to ours at 1 h (current study: 83.5%, Wright *et al.:* 80%).

Protocol 1 was shorter than protocol 2 making it a faster way to condition bees, while it also required no aversive US and less samples during training. However, protocol 1 did not result in the bees being able to discriminate between the novel infected and novel healthy samples. That indicates that they were not able to generalize between infected samples and to associate VOCs that commonly occur in infected samples with a reward. The bees correctly discriminated between the old infected and novel healthy samples which provided confirmation of the ability of the bees to recognise specific VOCs, but not to generalize over different infected samples. In contrast to protocol 1, protocol 2 resulted in a better discrimination ability between novel infected and healthy samples at both 1 h and 24 h retention, indicating that this protocol is more efficient for training bees for SARS-CoV-2 diagnostic testing. The differences between the two protocols outputs may be the consequence of the bees’ tendency to increase their attention during a learning task when faced with a potentially negative outcome (Avarguès-Weber *et al*., 2010; Chittka *et al*., 2003). In addition, both protocols differed in the number of conditioning rounds which could yield in a better memory. Finally, different samples were used in the conditioning phase of protocol 2, thus promoting generalization of response based on the common properties of all infected samples rather than on individual differences.

Conditioning of bees to SARS-CoV-2 derived VOCs could thus be further improved by focusing on the protocol that best worked in this study and add other elements that can make conditioning even more effective. Such an addition could be an extra training few hours after the original one or a different number of trials and alternative US. In our experiments we used appetitive-aversive conditioning due to the complexity and similarity of the odours the bees were trained for. In some cases, especially when bees are trained to fewer complex odours, the addition of a negative reinforcement can lead to lower discrimination and higher false positives (Aguiar *et al*., 2018).

Our results show that using single bees for diagnosis would have limited sensitivity and specificity, since the retention tests for protocol 2 showed that at 24 h post conditioning 67% of the bees correctly identified the infected sample (sensitivity) and 58% the healthy sample (specificity). A possible approach to improve diagnostic performance would be the use of multiple bees probing the same sample in parallel. In this case, diagnosis would be based on a defined number of bees (known as diagnostic threshold) reacting (expressing PER) to the sample being tested. We assessed such an approach by performing simulations where groups of 10 bees would be used to test a sample and identified that at least six bees would have to show PER for the sample to be considered positive. By taking this approach, the potential sensitivity of the test was predicted to be around 92% and the specificity around 86%. The current standard for laboratory diagnosis of active SARS-CoV-2 infection is the detection of viral RNA from respiratory specimens by real-time, reverse transcription polymerase chain reaction (qRT-PCR). Our predicted results on accuracy are comparable to the diagnostic performance of point of care (POC) tests such as RT-LAMP tests (without RNA extraction) and rapid antigen tests. These tests showed sensitivities higher than 70% for samples with Ct<33 or taken within the first week of symptom onset and specificities higher than 90% (Fowler *et al*.2021, Krüttgen *et al* 2021, Dinnes *et al*., 2021). In general, these POC tests require more than 10 minutes to produce a test result, whereas bees only require a few seconds to express PER (< 5 s). Dogs can also be trained to detect SARS-CoV-2 and provide results very quickly. However, dogs require much more time and resources to be trained compared with bees, and their sensitivity is lower than the honeybee test (dogs: sensitivity ranges from 65-82.6%; Eskandari *et al*., 2021; Jendrny *et al*., 2020). Moreover, dogs may be infected with SARS-CoV-2 whereas bees are not sensitive to the virus. In addition, bees can be employed in remote areas where microbiological laboratory facilities are not available. As such, it can be concluded that the honeybee test is a suitable alternative especially in situations where resources and laboratory equipment are scarce. This establishes the bee diagnostic test as an attractive monitoring method for developing countries and remote livestock communities, thanks to its low requirements and good diagnostic efficacy.

Our results suggest that honeybees could be used for SARS-CoV-2 diagnosis and could potentially be applied for diagnosis of other infectious diseases. Further research is needed in order to define the duration of their memory. It is clear that their memory is weaker 24 h after the experiments compared to 1 h after the training, which might be the result of complexity and similarity of the odours. We need to identify the crucial moment in time, in which their memory retention is compromised and further assess the performance with a wider range of Ct values. Here we only tested samples with a maximum Ct of 30 and given the limited number of samples tested, we cannot assume that the performance would be similar with higher Ct values. In addition, a formal diagnostic validation study is necessary to properly validate the diagnostic approach applied under field conditions and confirm the diagnostic potential predicted in this study. The diagnostic test proposed in this study has certain weaknesses that need to be improved. The need to use multiple bees in parallel along with the laborious process of conditioning bees manually can make the preparation of the test inefficient. In addition, the bees can only be used for testing a few samples before an extension of their memory would be observed due to the absence of reward during the tests. The bees will thus have to go through a few numbers of reactivating conditioning rounds before being again operational for testing.

## Conclusion

Our results indicate that the VOC profile differs between healthy and SARS-CoV2 infected minks and that honeybees can recognise these differences and discriminate between them. This performance suggests the presence of specific biomarkers, which could be explored by performing a Gas Chromatography/ Mass Spectrometry (GC/MS) analysis. Our experiments demonstrate that bees can effectively detect the presence of an infection in samples of an extensive range of Ct values. Once improved, a diagnostic test utilizing the learning abilities of honeybees might thus provide an important addition to the current monitoring system of zoonotic diseases in remote livestock farming systems.

## Acknowledgements

We thank Bregtje Smit for outstanding laboratory technical assistance. Dr. Hans Smid from the Laboratory of Entomology of Wageningen University, for his valuable insights and guidance, Saptarshi Mukhopadhyay from InsectSense B.V. and Edmund Taylor for their assistance and support throughout the project. This study was supported by the Wageningen University and Research KB37 research program and by funding from the European Union’s Horizon 2020 Research and Innovation programme under grant agreement No 773830: One Health European Joint Programme.”

## Competing interests

We declare no competing interests.

## Notes

### Competing Interest Statement

The authors have declared no competing interest.

## References

Aguiar, J. M. R. B. V. et al. (2018) ‘Can honey bees discriminate between floral-fragrance isomers?’, Journal of Experimental Biology, 221(14). doi: 10.1242/jeb.180844.

Avarguès-Weber, A., de Brito Sanchez, M.G., Giurfa, M. and Dyer, A.G., 2010. Aversive reinforcement improves visual discrimination learning in free-flying honeybees. PLoS One, 5(10), p.e15370.

Bates, D., Mächler, M., Bolker, B. and Walker, S., 2014. Fitting linear mixed-effects models using lme4. arXiv preprint arXiv:1406.5823.

Buljubasic, F. and Buchbauer, G. (2015) ‘The scent of human diseases: A review on specific volatile organic compounds as diagnostic biomarkers’, Flavour and Fragrance Journal, 30(1), pp. 5–25. doi: 10.1002/ffj.3219.

Chittka, L., Dyer, A.G., Bock, F. and Dornhaus, A., 2003. Bees trade off foraging speed for accuracy. Nature, 424(6947), pp.388–388.

Corman VM, Landt O, Kaiser M, Molenkamp R, Meijer A, Chu DK, et al. Detection of 2019 novel coronavirus (2019-nCoV) by real-time RT-PCR. Euro Surveill. 2020;25(3):2000045. 10.2807/1560-7917.ES.2020.25.3.2000045

Dinnes, Jacqueline, et al. “Rapid, point-of-care antigen and molecular-based tests for diagnosis of SARS-CoV-2 infection.” Cochrane Database of Systematic Reviews 3 (2021).

Eskandari, E., Marzaleh, M.A., Roudgari, H., Farahani, R.H., Nezami-Asl, A., Laripour, R., Aliyazdi, H., Moghaddam, A.D., Zibaseresht, R., Akbarialiabad, H. and Zoshk, M.Y., 2021. Sniffer dogs as a screening/diagnostic tool for COVID-19: a proof of concept study. BMC infectious diseases, 21(1), pp.1–8.

European Centre for Disease Prevention and Control. COVID-19 testing sequences and objectives. 15 September 2020. ECDC: Stockholm; 2020.

Fitzgerald, J. E. et al. (2017) ‘Artificial Nose Technology: Status and Prospects in Diagnostics’, Trends in Biotechnology. Elsevier Ltd, 35(1), pp. 33–42. doi: 10.1016/j.tibtech.2016.08.005.

Fowler, V.L., Armson, B., Gonzales, J.L., Wise, E.L., Howson, E.L., Vincent-Mistiaen, Z., Fouch, S., Maltby, C.J., Grippon, S., Munro, S. and Jones, L., 2021. A highly effective reverse-transcription loop-mediated isothermal amplification (RT-LAMP) assay for the rapid detection of SARS-CoV-2 infection. Journal of infection, 82(1), pp.117–125.

Gronenberg, W. et al. (2014) ‘Honeybees (Apis mellifera) learn to discriminate the smell of organic compounds from their respective deuterated isotopomers’, Proceedings of the Royal Society B: Biological Sciences, 281(1778). doi: 10.1098/rspb.2013.3089.

Hadagali, M. D. and Suan, C. L. (2017) ‘Advancement of sensitive sniffer bee technology’, TrAC - Trends in Analytical Chemistry. Elsevier Ltd, 97, pp. 153–158. doi: 10.1016/j.trac.2017.09.006.

Jendrny, P. et al. (2020) ‘Scent dog identification of samples from COVID-19 patients - A pilot study’, BMC Infectious Diseases. BMC Infectious Diseases, 20(1), pp. 1–7. doi: 10.1186/s12879-020-05281-3.

Kanazawa, M. et al. (2005) ‘Classical conditioned response of rectosigmoid motility and regional cerebral activity in humans’, Neurogastroenterology and Motility, 17(5), pp. 705–713. doi: 10.1111/j.1365-2982.2005.00691.x.

Krüttgen, A., Cornelissen, C.G., Dreher, M., Hornef, M.W., Imöhl, M. and Kleines, M., 2021. Comparison of the SARS-CoV-2 Rapid antigen test to the real star Sars-CoV-2 RT PCR kit. Journal of virological methods, 288, p.114024.

Matsumoto, Y. et al. (2010) ‘Revisiting olfactory classical conditioning of the proboscis extension response in honey bees: A step toward standardized procedures’, Journal of Neuroscience Methods. Elsevier B.V., 211(1), pp. 159–167. doi: 10.1016/j.jneumeth.2012.08.018.

Olsson, M. J. et al. (2014) ‘The Scent of Disease: Human Body Odor Contains an Early Chemosensory Cue of Sickness’, Psychological Science, 25(3), pp. 817–823. doi: 10.1177/0956797613515681.

Oreshkova, N., Molenaar, R.J., Vreman, S., Harders, F., Munnink, B.B.O., Hakze-van Der Honing, R.W., Gerhards, N., Tolsma, P., Bouwstra, R., Sikkema, R.S. and Tacken, M.G., 2020. SARS-CoV-2 infection in farmed minks, the Netherlands, April and May 2020. Eurosurveillance, 25(23), p.2001005.

Pavlov, I. P. (1927) ‘Conditioned reflexes: An investigation of the physiological activity of the cerebral cortex’, Annals of neurosciences, 17(3). doi: 10.5214/ans.0972-7531.1017309.

Probert, C.S., Khalid, T., Ahmed, I., Johnson, E., Smith, S. and Ratcliffe, N.M., 2009. Volatile organic compounds as diagnostic biomarkers in gastrointestinal and liver diseases. Journal of Gastrointestinal and Liver Disease, 18(3).

R.C., Team, 2013. R: A language and environment for statistical computing.

Robin, X., Turck, N., Hainard, A., Tiberti, N., Lisacek, F., Sanchez, J.C. and Müller, M., 2011. pROC: an open-source package for R and S+ to analyze and compare ROC curves. BMC bioinformatics, 12(1), pp.1–8.

Sethi, S., Nanda, R. and Chakraborty, T. (2013) ‘Clinical application of volatile organic compound analysis for detecting infectious diseases’, Clinical Microbiology Reviews, 26(3), pp. 462–475. doi: 10.1128/CMR.00020-13.

Shirasu, M. and Touhara, K. (2011) ‘The scent of disease: Volatile organic compounds of the human body related to disease and disorder’, Journal of Biochemistry, 150(3), pp. 257–266. doi: 10.1093/jb/mvr090.

Suckling, D. M. and Sagar, R. L. (2011) ‘Honeybees Apis mellifera can detect the scent of Mycobacterium tuberculosis’, Tuberculosis. Elsevier Ltd, 91(4), pp. 327–328. doi: 10.1016/j.tube.2011.04.008.

Sutherland, A. M. et al. (2010) ‘Classical conditioning of domestic honeybees to olfactory stimuli associated with grapevine powdery mildew infections’, pp. 90–92.

Sing, T., Sander, O., Beerenwinkel, N. and Lengauer, T., 2005. ROCR: visualizing classifier performance in R. Bioinformatics, 21(20), pp.3940–3941.

Takeda, K., 1961. Classical conditioned response in the honey bee. Journal of Insect Physiology, 6(3), pp.168–179.

Trabue, S. et al. (2010) ‘Speciation of volatile organic compounds from poultry production’, Atmospheric Environment, 44(29), pp. 3538–3546. doi: 10.1016/j.atmosenv.2010.06.009.

Wells, C.R., Townsend, J.P., Pandey, A. et al. Optimal COVID-19 quarantine and testing sequences. Nat Commun 12, 356 (2021). https://doi.org/10.1038/s41467-020-20742-8

Wilson, A. D. (2018) ‘Applications of electronic-nose technologies for noninvasive early detection of plant, animal and human diseases’, Chemosensors, 6(4), pp. 1–36. doi: 10.3390/chemosensors6040045.

Wright, G. A. et al. (2010) ‘Parallel reinforcement pathways for conditioned food aversions in the honeybee’, Current Biology, 20(24), pp. 2234–2240. doi: 10.1016/j.cub.2010.11.040.

